# Contributions of the gut microbiota and intestinal gluconeogenesis to metabolic adaptations in a mouse model of anorexia nervosa

**DOI:** 10.64898/2026.09.15.746633

**Authors:** Shiou-Ping Hviid-Chen, Zahra Boudra, Nadim Kassis, Soham Saha, Stéphanie Billon-Crossouard, Justine Vily-Petit, David Ribet, Pierre-Yves Barelle, Mikaël Croyal, Nicolas Ramoz, Virginie Tolle, Moïse Coëffier, Odile Viltart, Gilles Mithieux, Christophe Magnan, Céline Cruciani-Guglielmacci

## Abstract

Anorexia nervosa is a metabolic-psychiatric disorder characterized by severe food restriction, often accompanied by hyperactivity, associated with high mortality and the lack of specific pharmacological treatment. Despite a marked energy deficit, patients paradoxically maintain euglycemia, suggesting adaptations in energy homeostasis. This study aims to characterize the interactions between food restriction, the gut microbiota and peripheral organs in a mouse model mimicking anorexia nervosa.

Our results show that food-restricted mice exhibit improved glucose tolerance and increased expression of the gluconeogenic genes G6pc and Pck1 in the intestine, suggesting an adaptation in endogenous glucose production. Gut microbiota analysis reveals a marked shift in composition in food-restricted mice, with an increase in the *Lachnospiraceae* and *Marinifilaceae* families and a decrease in *Lactobacillaceae*. These changes are associated with metabolic parameters such as glycemia, body weight, and GLP-1 levels.

Transfer of microbiota from food-restricted mice into control mice improves glucose tolerance and increases the gluconeogenic gene expression. Furthermore, the improvement in glucose tolerance is abolished in mice lacking intestinal gluconeogenesis, suggesting that intestinal gluconeogenesis is required for the effects of the FR microbiota on glucose homeostasis.

Overall, these findings highlight a complex metabolic adaptation in a mouse model of anorexia nervosa, involving interactions between the gut microbiota, intestinal gluconeogenesis and maintenance of energy homeostasis.

## 1. INTRODUCTION

Anorexia nervosa (AN) is a metabolic-psychiatric disorder with the highest mortality rate among mental illnesses, characterized by severe food restriction, an intense fear of weight gain, and a distorted perception of body image (Auger et al. 2021; Kounatidis and Vallianou 2025). Despite decades of investigation in various disciplines like psychiatry, genetics and metabolism, the pathogenesis of AN is only partially understood and a specific pharmacological treatment is still missing. The main management of the AN currently relies on rehabilitation of normal nutrition and psychological interventions, with a high rate of relapse.

In AN patients, increased physical activity is often associated with food restriction, with 35–80% of patients who present excessive physical activity associated with a severe chronic food restriction as a result of their intense fear of gaining weight (Dalle Grave et al. 2008). More recently, the use of a continuous cardiac monitoring in patients suffering of acute AN evidences a negative correlation with minimal BMI and high level and moderate level physical activity (Duriez et al. 2021). Despite a marked energy deficit, patients paradoxically maintain normoglycemia or a stable subnormal glycemia (Pulini et al. 2025; Viltart et al. 2018), suggesting adaptations in energy homeostasis. The murine model of imposed chronic food restriction (FR), with or without voluntary access to an exercise wheel (FRW), recapitulates several key metabolic features of this disorder : maintained subnormal glycemia, increased glucose tolerance and insulin resistance (Méquinion et al. 2015). However, the underlying mechanisms allowing the long-term stabilization of body weight and maintenance of glucose homeostasis remain poorly understood.

Among them, the gut microbiota could be involved, as various changes in bacterial profiles have been measured in patients with AN (Di Lodovico et al. 2021), and the microbiota-gut-brain axis is well known to control whole body energy homeostasis (Wachsmuth et al. 2022). Interestingly, a dysbiosis is also observed in a mouse model of activity-based anorexia (ABA, which combines activity in a running wheel and time restricted food access), with in particular an increase in the abundance of *Clostridium cocleatum* and several *Lactobacillus* species, and a decrease in the abundance of *Burkholderiales* in ABA mice compared to control mice (Breton et al. 2021). Of note, most of the observed gut microbiota alterations are due to FR *per se* and are not affected by physical activity.

In the present study, we investigated the interplay between FR, the gut microbiota and peripheral organs in an AN-like mouse model, with the aim of elucidating the mechanisms underlying the maintenance of glucose homeostasis despite a marked energy deficit. To that end, we used a combination of the ABA and FRW mouse models: we imposed 50% restriction of food intake to isolated female mice and give them access to an exercise wheel. Indeed, in the original ABA protocol, developed by Routtenberg and Kuznezof in the 1960s, the rodents were isolated and did not survive more than 5-6 days (Routtenberg and Kuznesof 1967), whereas in the model named “food restriction and wheel” (FRW), female mice were housed in pairs (to avoid social stress and hypothermia) and exposed to 50% quantitative FR with free activity in a running wheel (Méquinion et al. 2015). The latter model allows to integrate both short term (15 days) and long term (10 weeks) protocols, permitting to study chronic metabolic adaptation. We have adapted this model by isolating the female mice during the FR protocol to ascertain the amount of food consumed, and we compensated for social stress and cold by providing housing enrichments. To note, this “modified FR/FRW model” is thus better adapted to gut microbiota transplantation studies.

In this work, we investigated the metabolic adaptations induced by chronic food restriction, focusing on glucose homeostasis and gut microbiota. We further examined whether the metabolic effects of the FR-associated microbiota involve intestinal gluconeogenesis, thereby exploring a potential microbiota– intestinal gluconeogenesis axis in the adaptation to chronic food restriction.

## 2. MATERIALS AND METHODS

### 2.1. Animals

#### Housing

Animal care and experimental procedures were approved by the Buffon Animal Experimentation Ethics Committee (CEEA-40) at Université Paris Cité, in accordance with French Ministry of Research regulations (APAFIS #40288-2022092811467226 v6). Mice were housed in a controlled environment with a temperature of 22 ± 1°C, a humidity of 50–70% and a 12-hour light/dark cycle (light on from 7 am to 7 pm). They were provided with standard chow diet (60.4% carbohydrate, 16.1% protein, 3.1% fat, 3.9% fiber; 2.79 kcal/g; Safe Diets-A04, Augy, France) and water unless otherwise specified. Each mouse was individually housed in ventilated cages and acclimated to daily handling.

#### Food restriction with/without wheel running protocol

Ten-week-old C57BL/6J female mice were obtained from Janvier Labs (Le Genest-Saint-Isle, France). On arrival, the mice underwent a one-week acclimation period. The following week, prior to the induction of the food restriction protocol, their baseline body weight and food intake were individually measured under *ad libitum* conditions. Mice were then randomly divided into four experimental groups. The AL (*ad libitum*) group mice served as the control group, where the mice had unrestricted access to a standard chow diet. The ALW (*ad libitum* with wheel access) group also had free access to food, but had a running wheel (12.4 cm in diameter; circumference, 39.0 cm) in their cage. AL and ALW mice had continuous access to food, whereas the daily food ration of FR and FRW mice was provided at 5 pm. In the FR (food restriction) group, the mice underwent a two-stage food reduction, a 30% reduction of their *ad libitum* food intake for three days, followed by a 50% reduction of their *ad libitum* food intake for three weeks. The FRW group (food restriction with running wheel access) followed the same food restriction but had voluntary access to a running wheel in their home cage. In accordance with the ethical approval, the body weights of the food-restricted mice were measured daily to ensure that they remained above 70% of their baseline body weight.

#### Intestine-specific G6pc knockout mice (G6pc^ΔVil^)

Mice were generated using a tamoxifen-inducible Cre/loxP system, as previously described (Penhoat et al. 2011). Briefly, G6pc lox/lox mice, in which exon 3 of the G6pc gene is flanked by loxP sites, were crossed with Villin-CreERT2 mice expressing the inducible CreERT2 recombinase under the control of the Villin promoter. Cre-mediated deletion of G6pc exon 3 was induced in adult mice (6-7 week-old) by tamoxifen administration (50mg/kg body weight; once daily for 5 consecutive days). Tamoxifen was prepared in 5% ethanol and 95% corn oil. Experiments were initiated 7 weeks after the final tamoxifen injection.

#### Animal euthanasia and tissue collection

On the day of euthanasia, food-restricted mice were provided with their food ration for 1 h in the morning, after which food was removed. All mice were then fasted for 6 h to ensure a consistent fasting duration across experimental groups. Mice were euthanized in the afternoon according to the approved experimental procedure. Following confirmation of death, tissues were rapidly collected and processed or stored as appropriate for analyses.

### 2.2. Metabolic phenotyping

#### Body weight and body composition

Mouse weight was measured using an electronic weighing device every day. Mouse body composition, including fat and lean mass (g), was measured precisely by the EchoMRITM-900 Analyzer.

#### Food intake

Food intake was measured daily at the same time each day. Under *ad libitum* conditions, the amount of food consumed was determined by weighing the remaining chow and calculating the difference from the amount provided on the previous day. Food intake was expressed as grams of chow consumed per mouse per day. These data were then used to calculate the amount of food provided to food-restricted mice, corresponding to 70% of ad libitum food intake for the first three days and 50% for the rest of the experiment.

For the metabolic tests, food-restricted mice were given access to their food ration for 1 h in the early morning, after which food was removed and mice were fasted for the duration specified in each experimental protocol. This procedure ensured that all mice underwent the same fasting duration before testing.

#### Oral glucose tolerance test

The oral glucose tolerance test (OGTT) was performed after three weeks of the food restriction protocol. Mice were fasted for five hours before receiving glucose (2 g/kg body weight; Glucose 30% B. Braun Medical) by oral gavage. Glycemia were measured using a glucometer (Glucofix Tech, Menarini Diagnostics, France) with a drop of blood sampled from the tail vein at baseline (0 minute) and at 15, 30, 45, 60, 90 and 120 minutes after glucose administration. 20 µL of blood was also collected at the baseline and at 15, 90 minutes after glucose gavage using capillary blood collection tube coated with heparin (Microvette® 300 Lithium heparin LH, France). Samples were immediately placed on ice. Plasma supernatants were collected by centrifugation of the blood samples at 4000 rpm for 10 minutes at 4°C. The collected plasma samples were then stored at −20°C until further measurements.

#### Intraperitoneal glucose tolerance test

The intraperitoneal glucose tolerance test (IPGTT) was performed following the same protocol as the OGTT, using the same glucose dose (2 g/kg body weight). The only difference was that glucose was administered by intraperitoneal (i.p.) injection rather than by oral gavage.

#### Oral glucose tolerance test with [14C]-2-deoxyglucose

Mice were fasted for five hours before the experiment and transferred to the experimental room before the beginning of the test. Blood glucose was first measured at baseline from the tail vein (T0). Mice were then weighed and received an oral glucose administration by gavage (2 g/kg body weight), immediately followed by an intraperitoneal injection of [^14^C]-2-deoxyglucose ([^14^C]-2DG, 5 µCi per mouse). Blood glucose was measured at 15, 30, 45, 60, 90 minutes after the gavage and the injection during the test. At each time point after [^14^C]-2DG injection, 10 µL of blood was collected from the tail vein and transferred into tubes containing 50 µL ZnSO₄. After homogenization, 50 µL Ba(OH)₂ was added, and samples were vortexed and centrifuged for 2 min at 10,000 rpm. Then, 50 µL of supernatant was transferred into scintillation vials and evaporated overnight. The dried pellets were resuspended in 500 µL distilled water, vortexed, mixed with 4.5 mL scintillation cocktail, and counted using a scintillation counter. At the end of the test, 90 min after injection, mice were euthanized by cervical dislocation immediately after the final blood collection. Tissues were rapidly collected, weighed, and placed in 500 µL 1N NaOH. Samples were digested for 1 h at 60°C until complete tissue dissolution. The homogenate was then processed in parallel with ZnSO₄/Ba(OH)₂ and perchloric acid 6% fractions to determine free [^14^C]-2DG and phosphorylated [^14^C]-2DG-6-phosphate. Radioactivity was measured by liquid scintillation counting. Tissue glucose uptake was calculated from the difference between total radioactivity measured in the perchloric acid fraction and free [^14^C]-2DG measured in the ZnSO₄ fraction.

#### Insulin tolerance test

The insulin tolerance test (ITT) was performed on day 20 of the food restriction protocol. Mice were fasted for five hours and then administered insulin (0.25 IU/kg body weight; NovoRapid Flexpen Insulin, Novo Nordisk, Sweden) by i.p. injection. Glycemia during the test was measured in the same way and at the same time points as the glucose tolerance test.

### 2.3. Gluconeogenic substrate tolerance tests

#### Pyruvate tolerance test

The pyruvate tolerance test (PTT) was performed to assess hepatic gluconeogenic capacity. Mice were fasted for 16 hours before receiving sodium pyruvate (2 g/kg body weight; 1.06619, Merck Millipore) by intraperitoneal (i.p.) injection. Blood glucose levels were measured using a glucometer (Glucofix Tech, Menarini Diagnostics) from tail vein blood samples collected at baseline (0 min) and at 15, 30, 45, 60, 90, and 120 min after pyruvate administration.

#### Glutamine tolerance test

The glutamine tolerance test was performed to evaluate gluconeogenesis from amino acid precursors. Mice were fasted for 6 hours before receiving L-glutamine (2 g/kg body weight; G3126, Sigma-Aldrich) by intraperitoneal (i.p.) injection. Blood glucose levels were measured using a glucometer (Glucofix Tech, Menarini Diagnostics) from tail vein blood samples collected at baseline (0 min) and at 15, 30, 45, 60, 90, and 120 min after glutamine administration.

### 2.4. Gastrointestinal motility, intestinal transit and intestinal permeability

Gastrointestinal motility was assessed using a carmine transit test. Mice were fasted for 16 h before the experiment and orally gavaged with a carmine solution containing 1.5% methylcellulose and 50 g/L carmine red at a volume of 20 µL/g body weight. Following gavage, mice were individually housed and monitored for the appearance of the first red-colored fecal pellet. Gastrointestinal transit time was defined as the time elapsed between gavage and the first appearance of carmine red in the feces.

Intestinal transit was assessed using a charcoal solution test. Mice were orally gavaged with 100 µL of a charcoal suspension (1 g charcoal in 5 mL H₂O) and sacrificed 1 h after gavage. The gastrointestinal tract was then collected, and the progression of the charcoal meal along the intestine was assessed.

Intestinal permeability was assessed in vivo using fluorescein isothiocyanate (FITC)-dextran. Mice were fasted for 6 h with ad libitum access to water and oral gavage with FITC-dextran (22 mg/mL in PBS; 20 mL/kg body weight). One hour after gavage, approximately 120 µL of blood was collected from the tail and centrifuged at 12,000 × g for 3 min at 4°C. Plasma was collected and diluted 1:3 (v/v) with PBS. Fluorescence was measured in a 96-well plate at excitation and emission wavelengths of 485 and 535 nm, respectively. Plasma FITC-dextran concentrations were determined using a standard curve prepared by diluting known concentrations of FITC-dextran in untreated plasma diluted 1:3 (v/v) with PBS.

### 2.5. Cecal microbiota transplantation

Recipient mice were pretreated with omeprazole (Inexium®, Grunenthal) and polyethylene glycol (Moviprep®, Norgine) prior to microbiota transplantation to facilitate donor microbiota engraftment. Donor cecal contents were diluted in sterile 0.9% NaCl to obtain a 20% (w/v) suspension, centrifuged at 3,000 × g for 3 min, and the supernatant was collected and maintained on ice until transplantation. Recipient mice received a daily oral gavage of omeprazole (200 µL/mouse) for three consecutive days. On day 4, mice were fasted for 1 h 30 min and received four oral gavages of polyethylene glycol solution (200 µL/mouse) at 20-min intervals. On day 5, mice received an additional oral gavage of omeprazole (200 µL/mouse), followed by oral administration of donor cecal suspension (100 µL/mouse). Thirty minutes later, mice were anesthetized with isoflurane and received a rectal flush with 150 µL sterile 0.9% NaCl, followed by rectal inoculation of 100 µL donor cecal suspension. Food was returned 1 h after transplantation. The cecal microbiota transplantation procedure was adapted from a previously published protocol (Péan et al., 2020), with modifications for the present mouse model. Cecal samples used for transplantation were collected from AL and FR donor mice after three weeks of food restriction. Recipient mice were then transplanted with microbiota from either AL (FMT-AL) or FR (FMT-FR) donors.

### 2.6. Microbiota composition

The microbiota composition was analyzed using 16S rRNA sequencing as previously described (doi:10.1016/j.clnu.2026.106681). Briefly, DNA from cecal contents was extracted using QIAamp Fast DNA Stool mini kit (Qiagen, Hilden, Germany), including a bead-beating step (0.1 mm zirconia silica beads). The University of Minnesota Genomics Center performed sequencing as follows: the V5-V6 region of the 16S rRNA gene was PCR-enriched using the primer pair V5F_Nextera (TCGTCGGCAGCGTCAGATGTGTATAAGAGACAGRGGATTAGATACCC) and V6R_Nextera (GTCTCGTGGGCTCGGAGATGTGTATAAGAGACAGCGACRRCCATGCANCACCT) and then underwent a library tailing PCR. The amplicons were purified, quantified and sequenced using an Element AVITI24 to produce 2 x 300 bp sequencing products. Then, the obtained sequences were denoised and chimera detected through DADA2 plugin of QIIME2 version 2025.7. The amplicon sequence variants (ASV) were associated to taxonomic identities using a pre-trained Naïve Bayes classifier with the SILVA v138.2 database and the plugin classify-sklearn. Alpha and beta diversity metrics were calculated with the QIIME2 diversity plugin. Data were normalized on the total number of ASV for differential analysis.

Taxonomic classification data and corresponding clinical metadata were processed in a Python environment using Pandas and NumPy. Bacterial taxonomy strings were parsed to isolate family-level annotations by searching for taxonomic family indicators (*f_ or _f*) or extracting the terminal clade identifier. Expression values for sample features sharing identical family classifications were aggregated and summarized by mean abundance. To ensure analytical integrity, sample identifier strings across the bacterial abundance matrix and metadata table were cleaned of leading and trailing whitespace and automatically aligned by sample matching. Metadata variables were restricted strictly to numeric features, and samples missing pairwise metadata or microbiome values were dropped pairwise to maintain valid statistical pairs. Pairwise association testing between numeric clinical metadata variables and bacterial family abundances was evaluated using non-parametric Spearman rank correlation (*ρ*). For each metadata–family pair, pairwise complete observations (N>3) were used to compute the correlation coefficient and its associated p-value via *scipy.stats.spearmanr*. To mitigate visual noise and focus on robust biological relationships, bacterial families were included in downstream visualization only if they demonstrated at least one statistically significant correlation (p < 0.05) across any metadata feature. Filtered correlation matrices were rendered as hierarchical heatmaps using Seaborn and Matplotlib. Diverging color scales (vlag) centered at zero were mapped to correlation coefficients ranging from −1 to +1. Statistical significance was visually annotated within each cell using standard threshold conventions: *, p<0.05; **, p<0.01; ***, p<0.001.

### 2.7. Metabolites prediction

Identification of metabolites affected by bacterial families were conducted using a large scale, grounded and targeted search on Seraj plaform (https://seraj.science). The research was conducted in February and March 2026 using an early-access to Seraj and its research-workflow functionality. PubMed and PubMed Central were used as literature sources, with search queries developed and refined iteratively. Seraj retrieved article abstracts and full texts where available and generated reports organized by bacterial taxon. The reports recorded metabolite–microorganism associations, their direction, evidence type, biological context, and citation identifiers. Seraj and human researchers checked the evidence; human checking of cited sources varied in extent. Before presenting the reports, Seraj performed internal quality checks, including citation removal and correction; a researcher subsequently approved the final association set before numerical conversion. Researchers converted all retained associations into the numerical species–metabolite interaction dataset without AI assistance for this conversion.

Interaction datasets mapping metabolites to individual microbial species were imported into a Python analysis environment along with network topological metrics. Missing interaction values were imputed as zero (representing neutral or unobserved interactions). Species-level identifiers were dynamically mapped to corresponding bacterial families. To incorporate network topology into functional associations, degree centrality metrics for each species were extracted from the network dataset. Species missing network centrality measures were assigned a default weight of 1.0. Raw species-level interaction scores were aggregated at the family level by summing species contributions within each taxonomic family to generate a metabolite-by-family raw interaction matrix. To identify latent functional associations and reduce sparsity in the raw interaction network, dimensionality reduction was performed on the aggregated metabolite-by-family matrix using Truncated Singular Value Decomposition (SVD) (*scikit-learn*). To account for family-level topological importance within the broader network, mean degree centrality scores were calculated for each bacterial family across its constituent species. These average family centrality scores were scaled via min-max normalization into a continuous weight factor bounded between 0.5 and 1.5 :

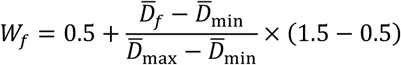

where *D̅_f_* represents the mean degree centrality of family *f*. The reconstructed latent interaction matrix was element-wise multiplied by these normalized family weights. Finally, weighted interaction values were mapped to continuous association probabilities bounded between 0 and 1 using a standard logistic sigmoid activation function :

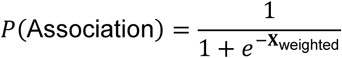

The resulting continuous probability matrix was melted into a tidy long-format structure to facilitate downstream data visualization and predictive statistical modeling.

### 2.8. Transcriptomic analysis

RNA sequencing was performed on liver and white adipose tissue samples from AL and FR mice. Mice were fasted for 6 h before tissue collection. RNA library preparation and sequencing were performed by Novogene (Cambridge, UK). Libraries were prepared following poly(A) enrichment, with approximately 25 million reads generated per sample.

Raw sequencing reads were quality-controlled using FastQC (v2.0.1) and Fastp (v0.20.0). Reads were aligned to the Mus musculus reference genome GRCm39 (mm39) using STAR (v2.7.10b). SAMtools (v1.13) and Qualimap (v2.2.2b) were used for alignment processing and quality assessment. Gene-level read counts were generated using featureCounts (v2.0.1), and MultiQC (v1.13) was used to summarize quality-control metrics. Differential gene expression analysis between AL and FR mice was performed using DESeq2. Gene Ontology (GO) enrichment analysis was performed in R using the clusterProfiler package, data visualization was performed using R. Kyoto Encyclopedia of Genes and Genomes (KEGG) pathway enrichment analysis was performed using Enrichr.

### 2.9. Statistical Analysis

Data are presented as means ± standard error of the mean (SEM). Normality was assessed using the Shapiro–Wilk test. Comparisons between two groups were performed using Student’s t-test (paired or unpaired, as appropriate), whereas comparisons involving more than two groups or experimental factors were performed using one- or two-way ANOVA, as appropriate. Repeated-measures ANOVA was used for experiments involving repeated measurements of the same animals. ANOVA was followed by Bonferroni-corrected post hoc multiple-comparison tests unless otherwise stated. The statistical test used for each experiment is specified in the corresponding figure legend. Differences were considered statistically significant at p < 0.05. Statistical analyses were performed using GraphPad Prism (GraphPad Software, La Jolla, CA, USA).

## 3. RESULTS

### 3.1. Increased glucose tolerance in an anorexia nervosa-like mouse model

We first aimed to validate that our modified AN-like mouse model using food restriction and wheel activity with one mouse per cage induced similar changes than the initial model (2 mice per cage, Mequinion et al., 2015) by measuring evolution of body weight, wheel activity, and body composition, including lean and fat mass (Figure 1A-E). Mice in the FR and FRW groups showed significant body weight loss, reaching approximately 20–25% after 10 days of the food-restriction protocol, and body weight then remained stable until the end of the protocol (Figure 1B). No difference was observed between the FR and FRW groups, indicating that wheel activity did not affect body weight loss. Although total wheel-running activity did not differ between the ALW and FRW groups (Figure 1C), their activity patterns differed, with FRW mice displaying more activity during the light phase and ALW mice predominantly running during the dark phase (data not shown). Body composition analysis showed a significant decrease in both lean mass and fat mass in FR and FRW mice, and FRW mice had lower fat mass than FR mice (Figure 1E).

**Figure 1.**
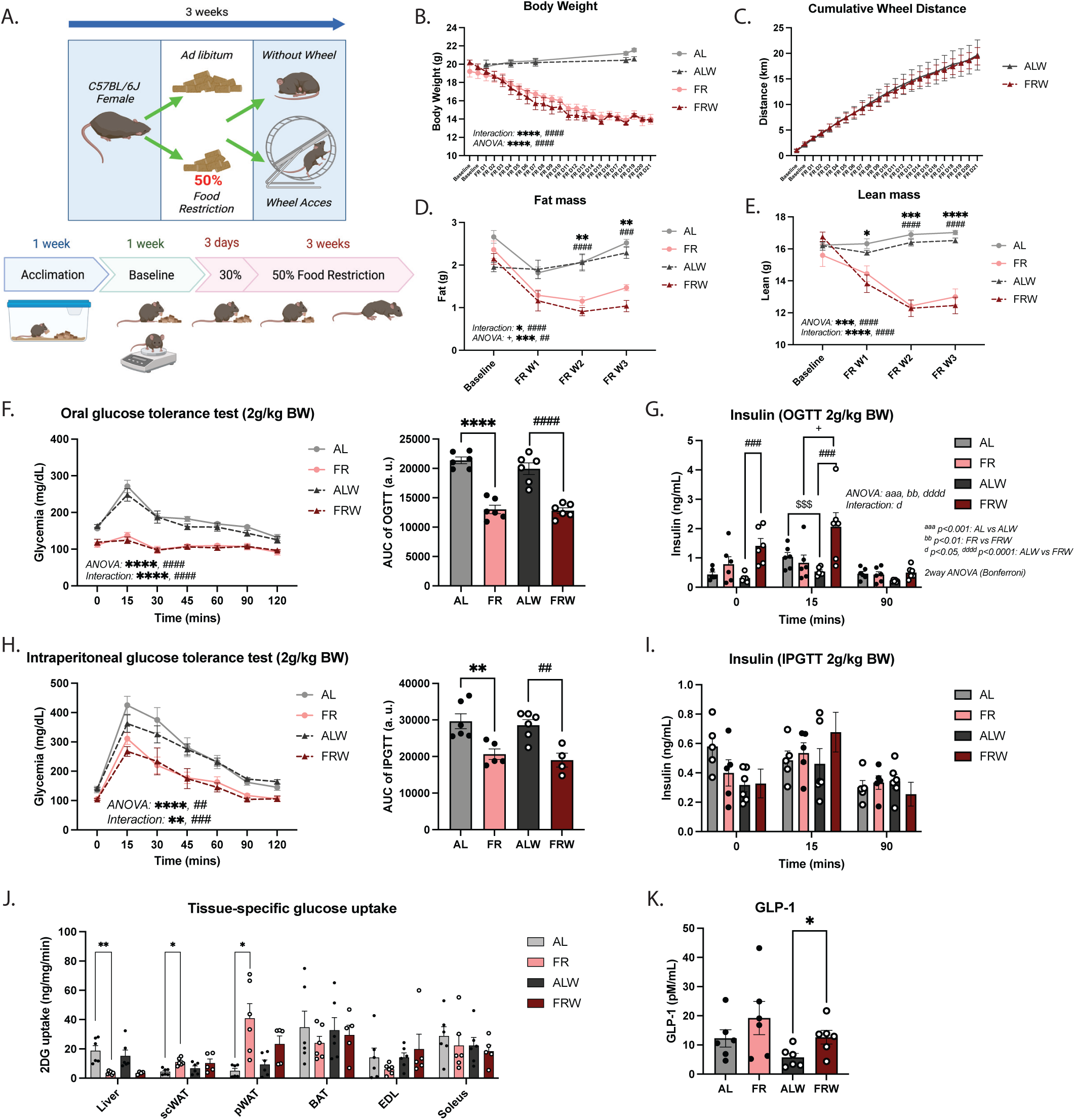
Metabolic adaptations induced by chronic food restriction in a mouse model of anorexia nervosa. A – Schematic illustration of the food restriction protocol used to mimic the metabolic phenotype of anorexia nervosa. B – Body weight (g) throughout the experimental protocol. C – Cumulative wheel activity (km) throughout the experimental protocol. D, E – Lean mass and fat mass measured throughout the experimental protocol. F – Left: oral glucose tolerance test (OGTT). Right: area under the curve (AUC). G – Plasma insulin levels during the OGTT. H – Left: intraperitoneal glucose tolerance test (IPGTT). Right: area under the curve (AUC). I – Plasma insulin levels during the IPGTT. J – 2-Deoxyglucose (2-DG) uptake in the gastrointestinal tract, liver, pancreas, skeletal muscle, brown adipose tissue (BAT), subcutaneous white adipose tissue (scWAT), and perigonadal white adipose tissue (pWAT). K – Plasma total GLP-1 levels. Data were obtained from female mice fed ad libitum (AL, grey), subjected to food restriction (FR, pink), fed ad libitum with wheel access (ALW, dark grey), or subjected to food restriction with wheel access (FRW, red). Data are presented as mean ± SEM (n = 6 mice per group). Statistical analyses are indicated in the corresponding panels. Unless otherwise indicated, data were analyzed using one-way ANOVA. OGTT and IPGTT data were analyzed using two-way ANOVA, whereas AUC values were analyzed using one-way ANOVA. Statistical significance is indicated as follows: *AL vs. FR, #ALW vs. FRW, $AL vs. ALW, and +FR vs. FRW. One, two, three, and four symbols indicate p < 0.05, p < 0.01, p < 0.001, and p < 0.0001, respectively.

We next aimed to determine the impact of food restriction and wheel activity on glucose tolerance. We performed an oral glucose tolerance test (OGTT) with measurement of insulinemia (Figure 1F–G). Improved glucose tolerance was observed in FR and FRW mice, as glycemia did not increase after oral glucose gavage (Figure 1F). In addition, FRW mice showed higher insulin levels before and 15 minutes after glucose gavage than FR mice (Figure 1G). FRW mice also exhibited insulin resistance in the insulin tolerance test (data not shown), suggesting that the improved glucose tolerance was partly due to increased insulin secretion during the OGTT. We then performed an intraperitoneal glucose tolerance test (IPGTT) to assess glucose tolerance independently of the gastrointestinal tract. FR and FRW mice still showed improved glucose tolerance compared with AL and ALW mice (Figure 1H), with no difference in insulin secretion (Figure 1I).

To further evaluate tissue-specific glucose uptake, we performed an OGTT combined with 2-deoxyglucose (2DG) and a 14C-radiolabeled tracer. This analysis revealed decreased glucose uptake in the liver and increased uptake in white adipose tissue in FR mice, with a similar trend toward decreased hepatic glucose uptake observed in FRW mice (Figure 1J). No difference was observed in brown adipose tissue or muscle (Figure 1J). Together, these results suggest that glucose uptake is altered in AN-like mice. To further explore the tissue-specific 2DG phenotype, we performed RNA-seq analysis in liver and adipose tissue. The transcriptomic changes are described in detail in section 3.6. Moreover, total plasma GLP-1 after five hours fasting was increased in FRW mice compared with ALW (Figure 1K). Overall, these findings suggest that the gastrointestinal tract may contribute to the improved glucose tolerance observed in this AN-like model.

### 3.2. Gastrointestinal changes and gluconeogenesis in AN-like mice

To better understand the involvement of the gastrointestinal tract in glucose regulation in the AN-like mouse model, we first measured intestine length, caecum weight, and stomach weight (Figure 2A– C). The results showed no difference in intestine length or caecum weight (Figure 2A–B), whereas increased stomach weight was observed in FR mice (Figure 2C). We next assessed intestinal barrier function by performing an intestinal permeability assay. Intestinal permeability was decreased in FR and ALW mice compared with AL mice (Figure 2D), suggesting enhanced intestinal barrier function in these groups. We also assessed gut transit time and gastric emptying; gut transit time was significantly decreased in FR and FRW mice (Figure 2E), whereas no difference was observed in gastric emptying time (data not shown). RT-qPCR revealed decreased mRNA expression of SGLT1, GLUT2, and GLUT5 in the jejunum in FR and FRW mice (Figure 2F–G), as well as decreased GLUT5 expression in the duodenum and ileum in FR mice (Figure 2H). Furthermore, we measured intestinal glucose absorption *ex vivo* in reversed intestinal segments from the duodenum, jejunum, and ileum, and found no difference in glucose absorption (data not shown). Taken together, these data suggest that glucose regulation is not directly altered by intestinal glucose absorption in FR and FRW mice.

**Figure 2.**
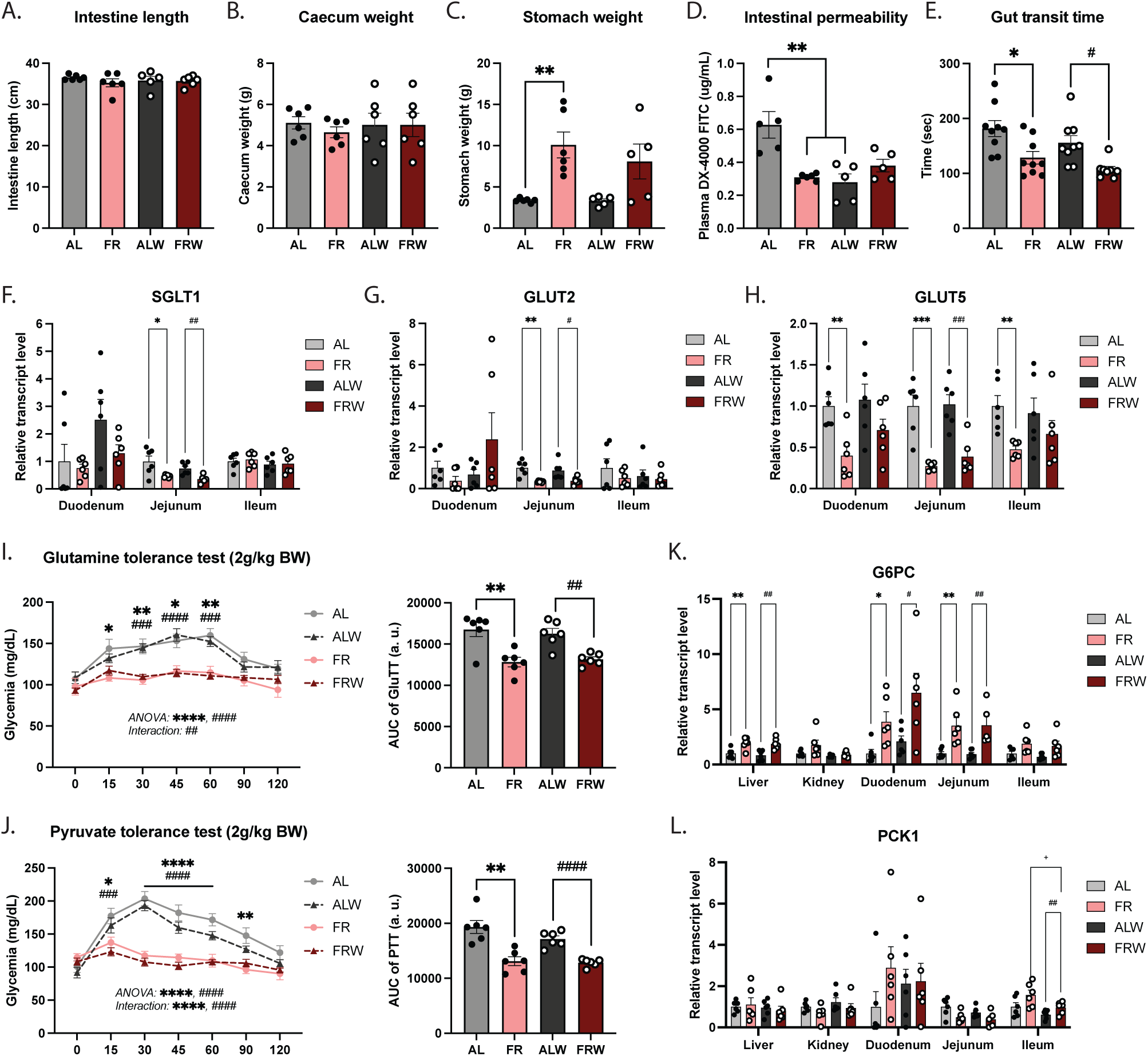
Intestinal adaptations to chronic food restriction. A – Intestine length (cm). B – Caecum weight (g). C – Stomach weight (g). D – Intestinal permeability assessed by FITC-dextran assay. E – Gastrointestinal transit time (s) assessed using the methylcellulose/dye transit assay. F, G, H – RT-qPCR analysis of the glucose transporter genes Slc5a1 (SGLT1) and Slc2a2 (GLUT2), as well as the fructose transporter gene Slc2a5 (GLUT5). I – Left: Glutamine tolerance test; Right: area under the curve (AUC) of the metabolic test. J – Left: Pyruvate tolerance test; Right: area under the curve (AUC) of the metabolic test. K, L – RT-qPCR analysis of the gluconeogenic genes G6pc and Pck1. Data were obtained from female mice fed ad libitum (AL, grey), subjected to food restriction (FR, pink), fed ad libitum with wheel access (ALW, dark grey), or subjected to food restriction with wheel access (FRW, red) (n = 6 per group). Data are presented as mean ± SEM. Statistical analyses are indicated in the corresponding panels and were performed using one-way ANOVA unless otherwise specified. Glutamine and pyruvate tolerance tests were analyzed using two-way ANOVA. Statistical significance is indicated as follows: *AL vs. FR, #ALW vs. FRW, $AL vs. ALW, and +FR vs. FRW. One, two, three, and four symbols indicate p < 0.05, p < 0.01, p < 0.001, and p < 0.0001, respectively.

We then evaluated gluconeogenesis by injecting gluconeogenic substrates, such as glutamine for renal and intestinal gluconeogenesis and pyruvate for hepatic gluconeogenesis. The results showed that glycemia in FR and FRW mice did not increase after glutamine or pyruvate injection (Figure 2I–J). Furthermore, we performed RT-qPCR on two key gluconeogenic enzymes, G6PC and PCK1. We found increased G6PC mRNA expression in the liver and intestine (Figure 2K), and increased PCK1 expression in the ileum in FR and FRW mice (Figure 2L). Overall, these findings suggest that gluconeogenesis may be altered in AN-like mice.

### 3.3. Food restriction alters gut microbiota composition

After observing differences between OGTT and IPGTT and no change in intestinal glucose absorption, we next evaluated whether food restriction altered the gut microbiota. We analyzed 16S rRNA gene sequences from cecal contents collected from AL, ALW, FR, and FRW mice. The minimum number of raw reads was approximately 120,000 per sample, indicating adequate sequencing depth for characterization of the bacterial community profiles present in the samples (Supplementary Figure 1). Alpha diversity at the genus level, assessed by Shannon index, showed no significant differences among the four groups (Figure 3A), indicating that within-sample diversity was not altered. By contrast, beta diversity using Bray-Curtis distance showed a significant difference between restricted groups (FR and FRW) and non-restricted groups (AL and ALW) (PERMANOVA analyses) (Figure 3B–C); similar results were observed with Jaccard distance (data not shown). These results suggest a major shift in gut microbiota composition after food restriction.

**Figure 3.**
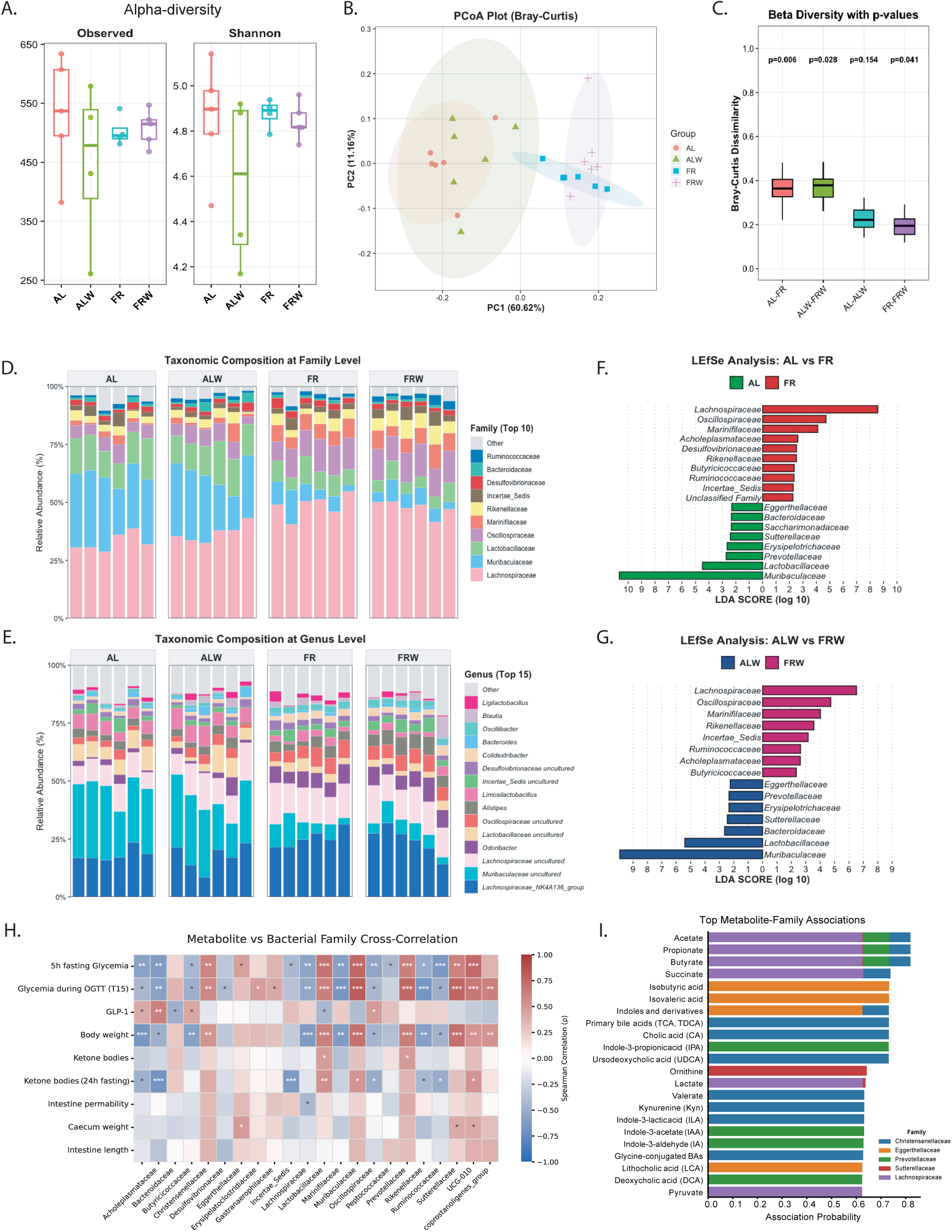
Chronic food restriction induces gut microbiota dysbiosis and alters microbiota– host metabolic associations. A – Alpha diversity of the gut microbiota assessed by the Observed and Shannon indices. B – Principal coordinates analysis (PCoA) based on Bray–Curtis distances showing β-diversity of the gut microbiota. C – Pairwise Bray–Curtis dissimilarity between experimental groups. D – Relative abundance of the ten most abundant bacterial families. E – Relative abundance of the fifteen most abundant bacterial genera. F – Linear discriminant analysis effect size (LEfSe) identifying differentially abundant bacterial families between AL and FR mice. G – LEfSe analysis identifying differentially abundant bacterial families between ALW and FRW mice. H – Cross-correlation analysis between bacterial families and metabolic parameters. I – Prediction of associations between microbial metabolites and bacterial families.

Taxonomic profiling at the family and genus levels, together with relative abundance bar plots and LEfSe analysis, revealed distinct microbial compositions between AL and FR mice as well as between ALW and FRW mice (Figure 3D–G), indicating that food restriction had a strong effect on microbiota composition, whereas wheel activity had a more limited effect. More specifically, *Lachnospiraceae* and *Oscillospiraceae* were significantly increased in food-restricted mice (Figure 3F–G), whereas *Muribaculaceae* and *Lactobacillaceae* were significantly decreased (Figure 3F–G).

To further investigate whether these microbial changes were related to glucose homeostasis, we performed a correlation analysis between the abundance of bacterial families and several metabolic parameters such as body weight or glycemia (Figure 3H). In addition, using a microbiota database, we predicted metabolites associated with bacterial families enriched in the food-restricted groups and found that short-chain fatty acids were highly associated with these families (Figure 3I). Collectively, these data indicate that food restriction strongly affects gut microbiota composition without altering alpha diversity, and that these microbial shifts may be linked to metabolic regulation, potentially through SCFA-related pathways.

### 3.4. Fecal microbiota transplantation from FR mice improves glucose tolerance and gluconeogenesis

To determine whether the microbiota from food-restricted mice contributes to glucose regulation, we performed cecal microbiota transplantation (FMT). Since wheel activity did not significantly alter gut microbiota composition, we foused on the FR group and assessed whether microbiota from AL or FR mice affected glucose metabolism in recipient mice pretreated with polyethylene glycol (PEG) for bowel cleansing (Figure 4A). Recipient mice receiving AL microbiota (FMT-AL) or FR microbiota (FMT-FR) were fed a normal chow diet without food restriction throughout the experiment.

**Figure 4.**
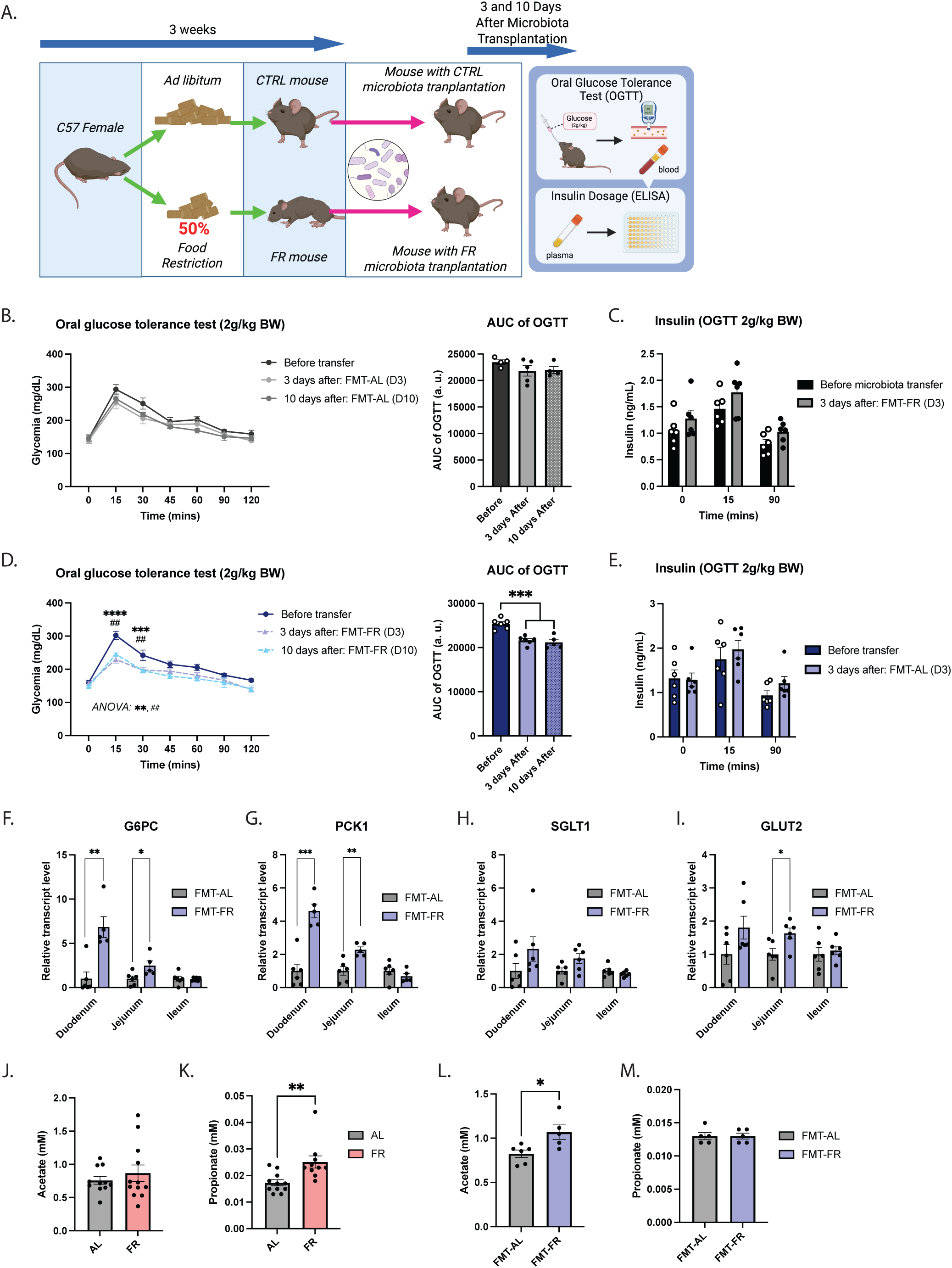
Metabolic modifications induced by cecal microbiota transplantation. A – Schematic illustration of the cecal microbiota transplantation protocol. B – Left: oral glucose tolerance test (OGTT) performed before and after cecal microbiota transplantation from control donors. Right: area under the curve (AUC). C – Plasma insulin levels during the OGTT before and after cecal microbiota transplantation from control donors. D – Left: OGTT performed before and after cecal microbiota transplantation from food-restricted donors. Right: area under the curve (AUC). E – Plasma insulin levels during the OGTT before and after cecal microbiota transplantation from food-restricted donors. F, G – RT-qPCR analysis of the gluconeogenic genes G6pc and Pck1. H, I – RT-qPCR analysis of the glucose transporter genes Slc5a1 (SGLT1) and Slc2a2 (GLUT2). J,K-Plasma Acetate and Propionate in AL and FR experimental groups. L, M – Plasma Acetate and propionate in mice following cecal microbiota transplantation. Data were obtained from female mice fed ad libitum (AL, grey), subjected to food restriction (FR, pink), and from female mice receiving microbiota from control (FMT-AL, grey) or food-restricted (FMT-FR, blue) donors. Data are presented as mean ± SEM (n = 6 mice per group). Statistical analyses are indicated in the corresponding panels. OGTT data were analyzed using two-way ANOVA, whereas AUC values and all other comparisons were analyzed using unpaired two-tailed Student’s t-test. Statistical significance is indicated by *p < 0.05, **p < 0.01, ***p < 0.001, and ****p < 0.0001.

Alpha diversity, assessed by the number of observed species and Shannon index, did not differ between the two recipient groups (Supplementary Figure 1). In contrast, beta diversity analysis using Bray-Curtis distance at the genus level showed a significant separation between the two groups based on PERMANOVA, indicating that the transferred microbiota differed between recipients (Supplementary Figure 1). Taxonomic profiling at the family and genus levels, together with relative abundance bar plots and LEfSe analysis, further revealed distinct microbiota compositions in the two recipient groups (Supplementary Figure 1).

Body weight was not affected by microbiota transfer (data not shown). OGTT was performed before transplantation and at 3 and 10 days after inoculation. Transfer of AL microbiota had no effect on glucose tolerance or insulinemia (Figure 4B–C). In contrast, mice receiving FR microbiota showed improved glucose tolerance without changes in insulinemia at both 3 and 10 days after inoculation (Figure 4D–E), suggesting that FR microbiota contributes to altered glucose regulation. Furthermore, pyruvate tolerance testing indicated a higher glycemic response in FMT-FR mice (data not shown). In addition, RT-qPCR showed increased expression of G6PC and PCK1 in FMT-FR mice (Figure 4F–G), suggesting enhanced gluconeogenesis. Finally, there was no difference in intestinal SGLT1 expression, whereas GLUT2 expression was increased in the jejunum (Figure 4H–I).

To better understand how the gut microbiota from food-restricted mice contributes to the regulation of glucose homeostasis, we next focused on a major class of microbiota-derived metabolites, the short-chain fatty acids (SCFAs). Gas chromatography–mass spectrometry (GC–MS) analysis revealed increased plasma propionate levels in food-restricted mice, and no difference in acetate levels (Figure 4J-K). Because the alterations in gut microbiota were primarily driven by food restriction rather than wheel access (Figure 3), the FR and FRW groups were pooled for this analysis. Furthermore, mice receiving microbiota from food-restricted donors (FMT-FR) exhibited increased plasma acetate levels compared with mice receiving microbiota from control donors (FMT-AL), no difference in plasma propionate level between two groups (Figure 4L-M). Collectively, these findings indicate that food restriction and transplantation of microbiota from food-restricted donors are associated with increased circulating SCFAs, suggesting that SCFAs may contribute to the improved glucose homeostasis observed in these mice.

### 3.5. Intestinal gluconeogenesis is required for the improved glucose tolerance induced by FR microbiota

To further investigate the involvement of intestinal gluconeogenesis in the improved glucose tolerance induced by the FR microbiota, we used an intestinal *G6pc* knockout mouse model (G6pc^ΔVil^). We therefore transferred AL or FR microbiota to control (littermate Cre−) or G6pc^ΔVil^ mice (Figure 5A). Before microbiota transplantation, glucose tolerance did not differ between groups (Figure 5B), indicating that intestinal *G6pc* deletion did not affect baseline glucose tolerance under these conditions. At 3 and 10 days after microbiota transplantation, control mice receiving FR microbiota showed improved glucose tolerance, consistent with our previous findings (Figure 5C and 5D). In contrast, this improvement was not observed in G6pc^ΔVil^ mice receiving FR microbiota (Figure 5C and 5D). Together, these results suggest that the improvement in glucose tolerance induced by the FR microbiota depends on intestinal gluconeogenesis.

**Figure 5.**
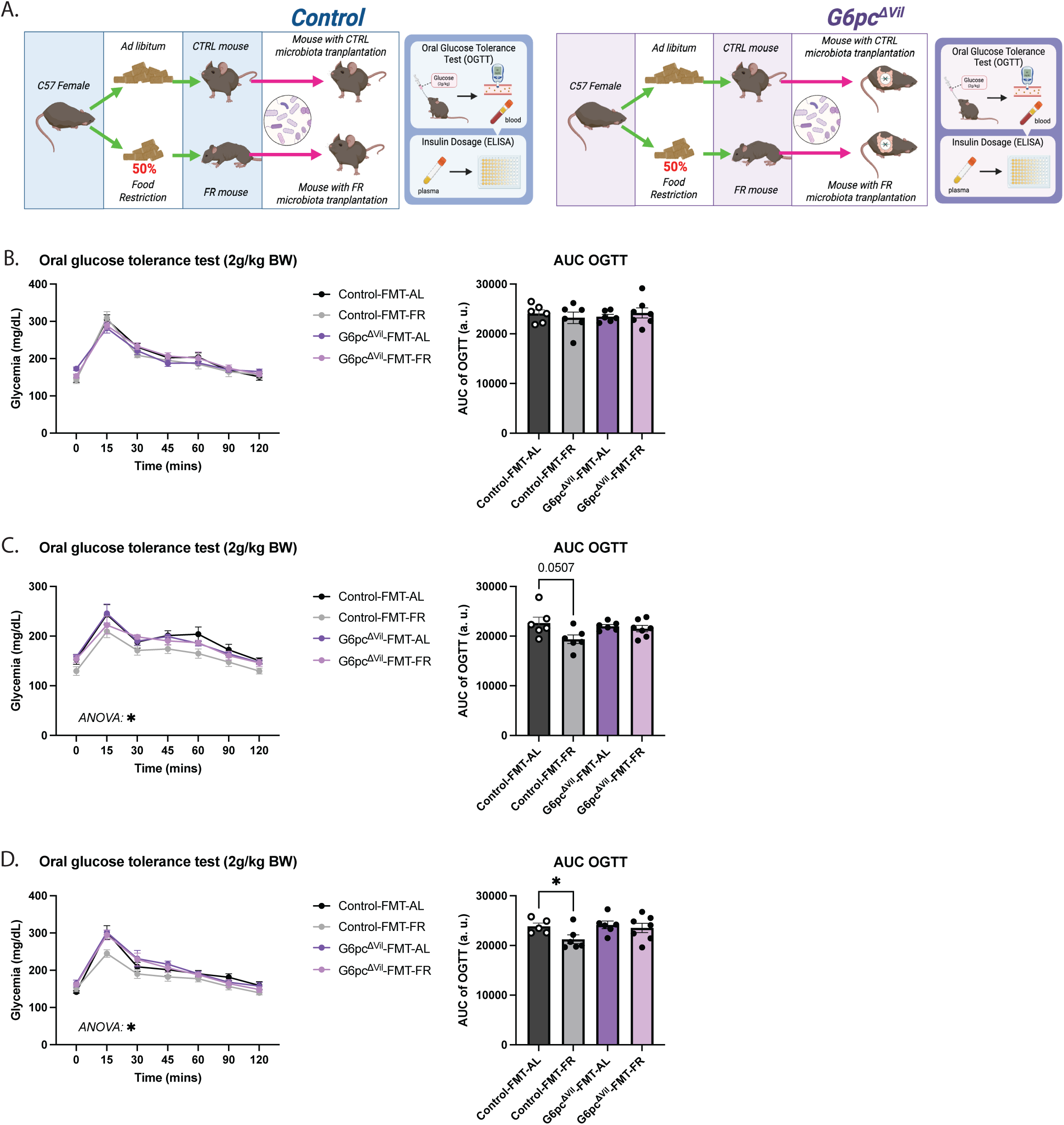
Glucose tolerance improvement following microbiota transplantation requires intestinal gluconeogenesis. A – Schematic illustration of the cecal microbiota transplantation protocol in Control (left) and G6pc^ΔVil^ mice (right). B – Left: oral glucose tolerance test (OGTT) performed before cecal microbiota transplantation from control AL or FR donors. Right: area under the curve (AUC). C - Left: oral glucose tolerance test (OGTT) performed 3 days after cecal microbiota transplantation from control AL or FR donors. D - Left: oral glucose tolerance test (OGTT) performed 10 days after cecal microbiota transplantation from control AL or FR donors. Data are presented as mean ± SEM (n = 6 mice per group). Statistical analyses are indicated in the corresponding panels. OGTT data were analyzed using two-way ANOVA, whereas AUC values and all other comparisons were analyzed using unpaired two-tailed Student’s t-test. Statistical significance is indicated by *p < 0.05, **p < 0.01, ***p < 0.001, and ****p < 0.0001.

### 3.6. Chronic food restriction profoundly changes liver and WAT transcriptomes

To investigate the transcriptional adaptations induced by chronic food restriction, RNA sequencing was performed on liver tissue from AL and FR mice. Principal component analysis (PCA) revealed a clear separation between AL and FR mice, indicating distinct hepatic transcriptional profiles between the two groups (Supplementary Figure 2A). Consistently, the volcano plot identified numerous differentially expressed genes (DEGs) in FR mice compared with AL controls (Supplementary Figure 2B). The heatmap of the 50 most differentially expressed genes further confirmed the distinct gene expression patterns between the two groups (Supplementary Figure 2C). GO enrichment analysis revealed that genes upregulated in FR mice were primarily associated with membrane transport and ion transport, including metal ion and sodium ion transmembrane transport, as well as solute sodium symporter activity (Supplementary Figure 2D). Immune-related processes, including MHC class II protein complex assembly and antigen presentation, were also significantly enriched among the upregulated genes (Supplementary Figure 2D). In contrast, downregulated genes were significantly enriched in lipid metabolic pathways, including fatty acid metabolism, steroid and sterol biosynthesis, isoprenoid metabolism, and peptide transport (Suppl. Fig. 2E).

Consistently, KEGG pathway enrichment showed that upregulated genes were associated with immune-related pathways, including primary immunodeficiency (Suppl. Fig. 2F), and that downregulated genes were enriched in pathways related to terpenoid backbone biosynthesis, retinol metabolism, galactose metabolism, starch and sucrose metabolism, adipocytokine signaling, and insulin resistance (Suppl. Fig. 2G). Overall, these results indicate that chronic food restriction profoundly changes the hepatic transcriptome, promoting coordinated changes in gene expression that reduce metabolic processes while increasing the expression of genes involved in membrane transport and immune-related processes.

RNA sequencing was also conducted on subcutaneous WAT from AL and FR mice. PCA showed a distinct separation between the two groups, confirming the different transcriptional profiles (Suppl. Fig. 3A). Furthermore, the volcano plot analysis identified a large number of DEGs in FR mice relative to AL controls (Suppl. Fig. 3B). In addition, the heatmap of the top 50 most significantly altered genes further distinguished the two experimental groups (Suppl. Fig. 3C).

GO enrichment analysis revealed that genes upregulated under food restriction were enriched in immune-related biological processes, including leukocyte-mediated immunity, lymphocyte activation, adaptive immune responses, and cytokine signaling (Suppl. Fig. 3D). Conversely, downregulated genes were enriched in lipid and sterol metabolic pathways, including fatty acid metabolism, cholesterol biosynthesis, steroid metabolism, and acetyl-CoA pathways (Suppl. Fig. 3E). KEGG pathway enrichment analysis confirmed these findings (Suppl. Fig. 3F and 3 G).

Together, these results demonstrate that chronic food restriction modulates the transcriptome of subcutaneous WAT, resulting in coordinated changes in immune- and metabolism-related pathways.

## 4. DISCUSSION

To investigate the metabolic adaptations associated with AN, we adapted the food restriction with wheel activity (FRW) mouse model previously described by Méquinion et al. (Méquinion et al. 2015). In the original protocol, mice were housed in pairs; however, group housing may influence both food consumption and wheel-running behavior. To allow accurate monitoring of individual food intake and physical activity, mice were therefore individually housed in the present study. Our results showed stable glycemia and improved glucose tolerance in both FR and FRW mice despite the marked body weight loss induced by chronic food restriction, suggesting that mice develop metabolic adaptations enabling them to maintain physiological function during prolonged energy deficiency. Comparable adaptations have been reported in patients with AN, who often maintain relatively stable glycemia despite severe and prolonged undernutrition (Pulini et al. 2026). This is particularly notable because glucose availability would be expected to decline as nutrient intake and energy stores become progressively depleted. Preserving glucose homeostasis is essential for sustaining glucose-dependent organs, particularly the brain and thus represents a critical survival adaptation during starvation (Mergenthaler et al. 2013). These findings underline the metabolic flexibility of the organism under extreme nutritional conditions and point to coordinated responses across multiple tissues and physiological systems rather than a single organ or pathway, which we discuss in the following sections. Interestingly, although excessive physical activity is considered a hallmark of AN, wheel-running activity had relatively limited effects on most of the metabolic and microbial parameters investigated in the present study. In contrast, food restriction consistently emerged as the primary driver of the observed adaptations. These findings suggest that chronic energy deficiency may play a more prominent role than physical activity in shaping the metabolic phenotype associated with AN.

One of the most important findings of this study is that chronic food restriction profoundly altered gut microbiota composition and that these microbial changes appear to be involved in the regulation of glucose homeostasis. Gut microbiota dysbiosis has been consistently reported in patients with AN as well as in ABA mouse models (Breton et al. 2021; Morita et al. 2015). Likewise, the contribution of the gut microbiota to glucose metabolism has been well documented (Kasubuchi et al. 2015; Portincasa et al. 2022). However, to our knowledge, few studies have investigated whether the microbiota alterations observed in AN contribute to the metabolic adaptations associated with the disease. Interestingly, genetic studies have reported negative genetic correlations between AN and several metabolic disorders, including obesity and type 2 diabetes (Watson et al. 2019). These observations raise the question of whether some physiological adaptations associated with AN may confer protection against metabolic dysfunction. In the present study, transplantation of microbiota from food-restricted mice improved glucose tolerance in recipient mice, suggesting that the AN-associated microbial profile contributes, at least in part, to the regulation of glucose metabolism. Importantly, although this phenotype may appear metabolically beneficial in the context of obesity or type 2 diabetes, it should not necessarily be considered beneficial in AN. Indeed, adaptive mechanisms that promote energy conservation and facilitate food restriction may contribute to the maintenance of the disease. In support of this idea, recent studies have suggested that gut microbiota alterations in AN may influence behavioral traits associated with the disorder (de Clercq et al. 2019; Kleiman et al. 2015; Fan et al. 2023). Therefore, the microbiota changes observed in AN may simultaneously contribute to metabolic adaptations and behavioral alterations.

We observed that transplantation of the microbiota from FR mice improved glucose homeostasis in recipient mice; however, the microbial composition of FMT-FR recipients was not consistently identical to that of the FR donor mice. Although the same bacterial families were not enriched in both groups, FMT-FR recipients nevertheless exhibited a distinct microbial profile compared with FMT-AL recipients. This finding is consistent with previous studies showing that fecal microbiota transplantation does not completely reproduce the donor microbiota in recipients, but can still confer beneficial physiological effects (Kowalska et al. 2026; Staley et al. 2016). Similarly, in patients with AN, FMT has been reported to improve clinical and metabolic outcomes despite incomplete engraftment of the donor microbiota (de Clercq et al. 2019). Interestingly, the increased expression of gluconeogenic genes (G6pc and Pck1) together with elevated plasma SCFA concentrations is consistent with a study published in 2014 demonstrating that SCFAs produced by the gut microbiota from a fiber-enriched diet improve metabolic regulation through the activation of intestinal gluconeogenesis (De Vadder et al. 2014). In addition, intestinal gluconeogenesis is known to be induced during fasting and by high-protein diets. Beyond its role in glucose homeostasis, intestinal gluconeogenesis has also been associated with increased satiety and reduced anxiety through gut-brain neural circuits (Penhoat et al. 2011; Sinet et al. 2021). Therefore, the increase in plasma SCFAs and gluconeogenic gene expression observed in our model raises the possibility that intestinal gluconeogenesis contributes to the adaptations induced by chronic food restriction. This hypothesis is particularly relevant in the context of AN, as patients often consume diets enriched in fiber and protein (Nova et al. 2001; Pettersson et al. 2021). Importantly, our results provide direct evidence that intestinal gluconeogenesis is required for the regulation of glucose homeostasis by the FR-associated microbiota. Indeed, while transplantation of FR microbiota improved glucose tolerance in control mice, this effect was abolished in mice lacking intestinal G6pc, demonstrating that the metabolic effects of the FR microbiota depend on intestinal gluconeogenesis. We therefore propose that chronic starvation induces gut microbiota dysbiosis, leading to altered SCFA production, which may stimulate intestinal gluconeogenesis. Through gut-brain communication, this pathway could contribute not only to the maintenance of glucose homeostasis but also to behavioral adaptations such as increased satiety and reduced perception of hunger. Although the involvement of SCFAs in this pathway remains to be confirmed, this mechanism could partly explain how AN patients maintain severe food restriction despite chronic undernutrition.

Beyond metabolic regulation, chronic food restriction may also influence immune system. Patients with AN have been reported to be relatively protected from the infection since the 1970s (Tk and E 1978). Several mechanisms have been proposed to explain this apparent paradox. For example, intestinal permeability has been reported to be reduced in AN patients (Monteleone et al. 2004), which is consistent with our observations in food-restricted mice. A reduced intestinal permeability may limit the translocation of pathogens and bacterial products across the intestinal barrier, thereby contributing to the preservation of immune function (Fine et al. 2019; Ghosh et al. 2020). In addition, several studies have reported alterations in immune responses in AN patients, including increased T-cell responsiveness and changes in B-lymphocyte populations (Elegido et al. 2017; Gibson and Mehler 2019). Interestingly, pathway enrichment analysis performed in white adipose tissue from our food-restricted mice also revealed upregulated immune-related pathways, suggesting that chronic food restriction induces specific immunological adaptations rather than a generalized immune suppression. However, not all immune alterations appear to be regulated in the same direction. For example, decreased neutrophil adherence has been reported in AN patients (Gibson and Mehler 2019), whereas pathways associated with leukocytes adhesion were upregulated in the white adipose tissue of FR mice. These apparent discrepancies may reflect tissue-specific adaptations or differences between human AN and experimental models of chronic food restriction.

One of the intriguing findings of this study is that glucose uptake was increased in white adipose tissue despite chronic food restriction and marked body weight loss after a glucose challenge. This observation may appear paradoxical, as adipose tissue is generally considered an energy storage organ that would be expected to be reduced during prolonged undernutrition. However, a similar phenomenon has been reported in patients with anorexia nervosa during nutritional rehabilitation. During weight recovery, AN patients preferentially accumulate adipose tissue, particularly visceral and intramuscular fat depots, compared with weight-matched healthy controls (Grinspoon et al. 2001; Mayer et al. 2005). For subcutaneous adipose tissue, the literature is less consistent, with some studies reporting increased fat accumulation after weight restoration whereas others found no significant differences (Mayer et al. 2005; Zamboni et al. 1997). Taken together, these observations suggest that chronic food restriction induces long-lasting metabolic adaptations that favor the restoration and preservation of energy stores. Increased glucose uptake by adipose tissue may therefore represent a preparatory mechanism aimed at restoring lipid reserves when nutrients become available. From an evolutionary perspective, such adaptations may increase the ability of the organism to survive with future periods of food deficiency. In addition, the restoration of adipose tissue may be required to recover endocrine functions that depend on fat mass, including the production of adipose-derived hormones and the normalization of reproductive function. Therefore, the preferential utilization of glucose by adipose tissue may reflect a coordinated adaptive response to chronic energy deficit rather than a maladaptive phenomenon.

In conclusion, we provide further evidence supporting the concept that AN is a metabo-psychiatric disorder. We observed multiple alterations, including dysbiosis, metabolic changes and adaptations in peripheral organs such as the liver and adipose tissue. Together, these findings suggest that chronic food restriction induces coordinated adaptations through the peripheral organs and the intestinal microbiota. Importantly, these adaptations may not simply represent compensatory responses to chronic energy deficit. Instead, some of them may actively contribute to the maintenance and reinforcement of anorexic behaviors, thereby establishing a self-sustaining vicious cycle. At the same time, several of the mechanisms identified in this work, such as the improvement in glucose tolerance mediated by the intestinal microbiota, may have broader implications beyond AN and could inspire novel therapeutic approaches for metabolic disorders such as obesity and type 2 diabetes.

## SUPPLEMENTAL FIGURES LEGENDS

**Suppl. Fig. 1.**
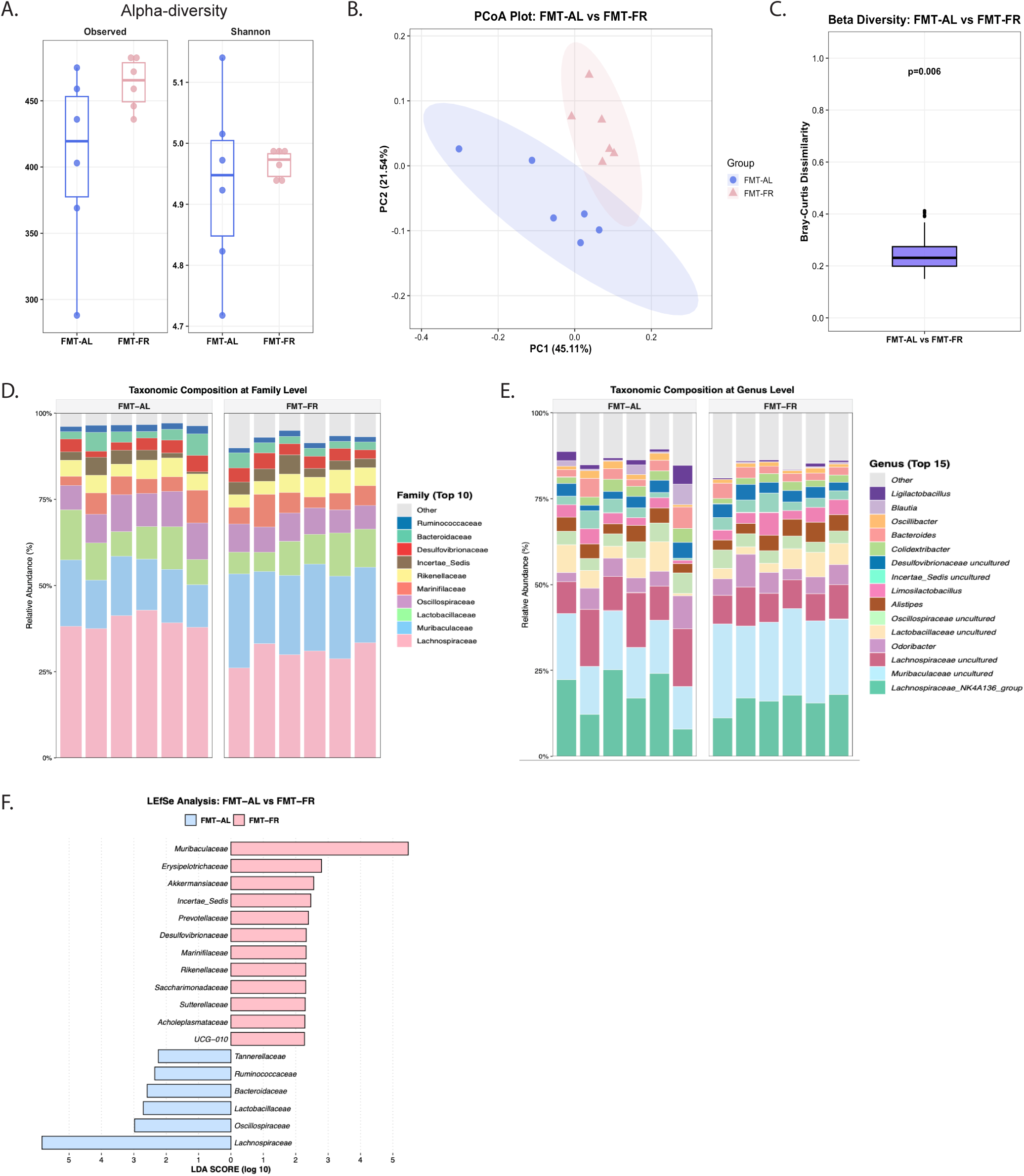
Changes in the gut microbiota following cecal microbiota transplantation. A – Alpha diversity of the gut microbiota assessed by the Observed and Shannon indices. B – Principal coordinates analysis (PCoA) based on Bray–Curtis distances showing β-diversity of the gut microbiota. C – Pairwise Bray–Curtis dissimilarity between experimental groups. D – Relative abundance of the ten most abundant bacterial families. E – Relative abundance of the fifteen most abundant bacterial genera. F – Linear discriminant analysis effect size (LEfSe) identifying differentially abundant bacterial families between FMT-AL and FMT-FR mice.

**Suppl. Fig. 2.**
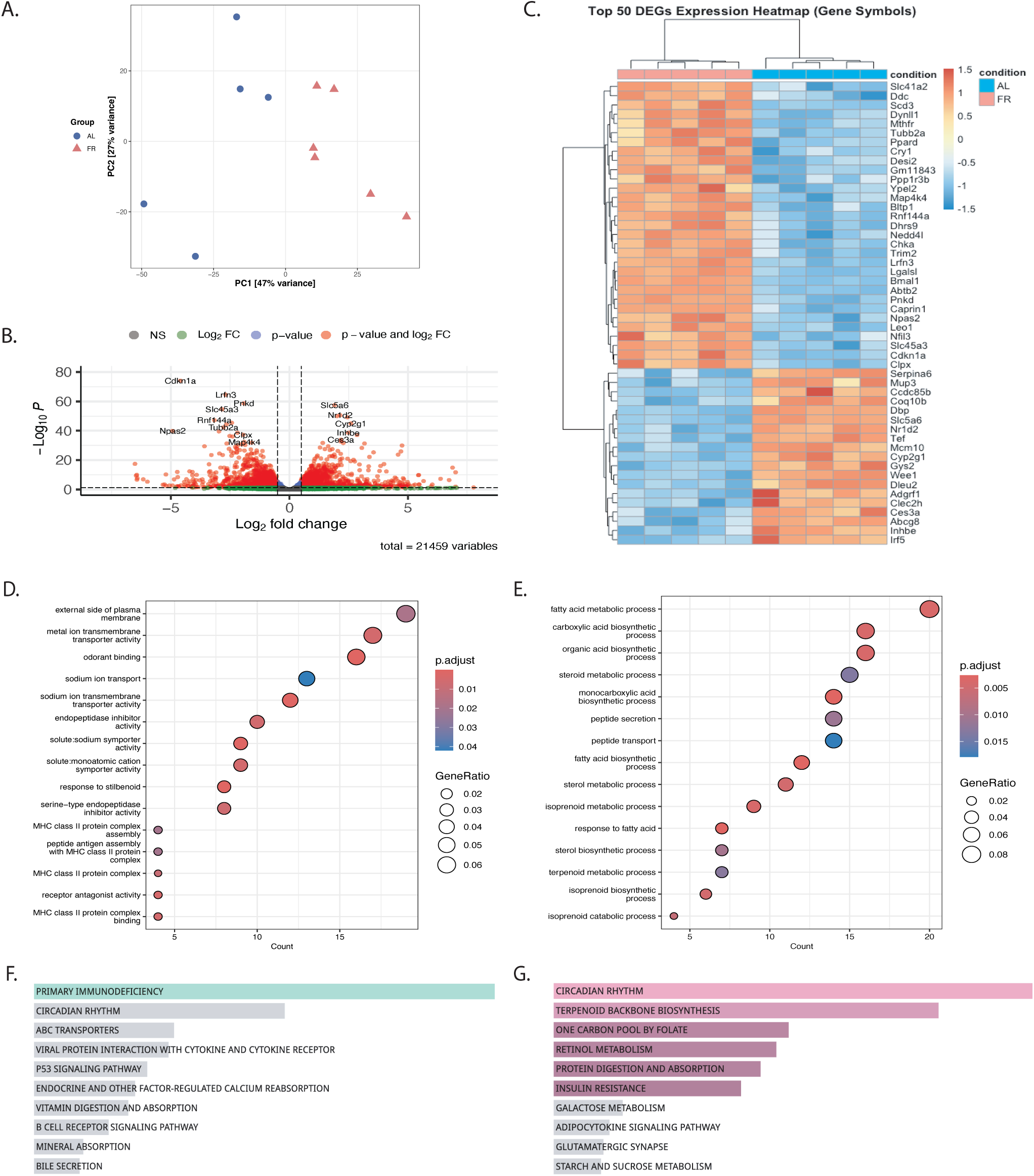
Chronic food restriction induces transcriptional modulations of the liver. A – Principal component analysis (PCA) of liver RNA-seq samples from AL and FR mice. B – Volcano plot showing differentially expressed genes between AL and FR mice. C – Heatmap of the 50 most differentially expressed genes identified by RNA sequencing. D – Gene Ontology (GO) enrichment analysis of upregulated genes. E – GO enrichment analysis of downregulated genes. F – Kyoto Encyclopedia of Genes and Genomes (KEGG) pathway enrichment analysis of upregulated genes. G – KEGG pathway enrichment analysis of downregulated genes. Data were obtained from liver tissue of female mice fed ad libitum (AL) or subjected to food restriction (FR). Principal component analysis was based on variance-stabilized transformed counts. Differential gene expression analysis was performed using DESeq2. Gene Ontology (GO) and Kyoto Encyclopedia of Genes and Genomes (KEGG) enrichment analyses were performed on significantly differentially expressed genes.

**Suppl. Fig. 3.**
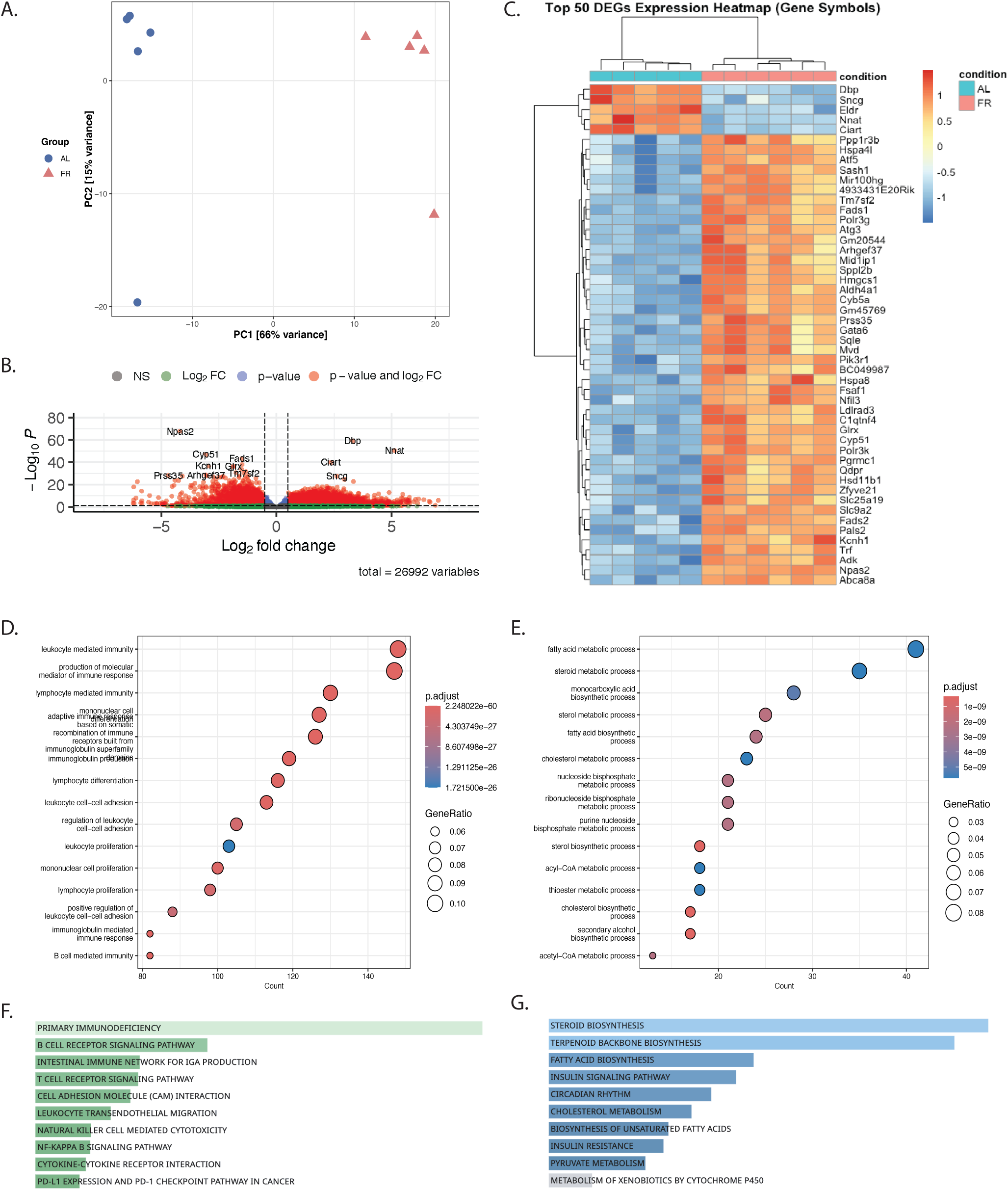
Chronic food restriction induces transcriptional modulations of the subcutaneous white adipose tissue. A – Principal component analysis (PCA) of subcutaneous white adipose tissue RNA-seq samples from AL and FR mice. B – Volcano plot showing differentially expressed genes between AL and FR mice. C – Heatmap of the 50 most differentially expressed genes identified by RNA sequencing. D – Gene Ontology (GO) enrichment analysis of upregulated genes. E – GO enrichment analysis of downregulated genes. F – Kyoto Encyclopedia of Genes and Genomes (KEGG) pathway enrichment analysis of upregulated genes. G – KEGG pathway enrichment analysis of downregulated genes. Data were obtained from white adipose tissue of female mice fed ad libitum (AL) or subjected to food restriction (FR). Principal component analysis was based on variance-stabilized transformed counts. Differential gene expression analysis was performed using DESeq2. Gene Ontology (GO) and Kyoto Encyclopedia of Genes and Genomes (KEGG) enrichment analyses were performed on significantly differentially expressed genes.

